# Multi-Transition Systems: A theory for neural spatial navigation

**DOI:** 10.1101/174946

**Authors:** Nicolai Waniek

## Abstract

Spatial navigation is considered fundamental for animals and is attributed primarily to place and grid cells in the rodent brain. Commonly believed to either perform path integration or localization, the true objective of grid cells, their hexagonal grid fields, and especially their discrete scales remain puzzling. Here it is proposed that grid cells efficiently encode transitions in sequences. A biologically plausible model for dendritic computation in grid cells is presented. A network of competitive cells shows positive gridness scores early in simulations and realigns the orientation of all cells over time. Then, a scale-space model of grid cells is introduced. It improves behaviorally questionable run-times of a single scale significantly by look-ahead in multiple scales, and it is shown that the optimal scale-increment between consecutive scales is *√*2. Finally, a formal theory for sequences and transitions is stated. It is demonstrated that hexagonal transition encoders are optimal to encode transitions in Euclidean space and emerge due to the sampling theorem. The paper concludes with a discussion about the suggested purpose, makes testable predictions, and highlights relevant connections to computational neuroscience as well as computer science and robotics.

## I. INTRODUCTION

Decades of research uncovered neurons which are relevant for navigation and express correlation with spatial information. The probably most prominent ones are Head Direction (HD) cells, Place Cells (PCs), and Grid Cells (GCs) [1], all of which can be found in the Hippocampal Formation (HF) [2]. Their combined representations are thought to form a map of the surrounding environment [1], [3], just as anticipated by Edward Tolman in his proposal of a *cognitive map* in 1948 [4].

HD cells show preferential tuning towards directions [5]–[7]. The firing rate of an HD cell is maximal when the animal faces the neuron’s preferred direction [8], [9]. Thus, a network of HD cells can be considered to resemble a compass [10].

A PC is active only when an animal is in one particular or a few randomly distributed locations of the environment, called place fields [11]–[14]. PCs are pyramidal neurons and were discovered in the areas Cornu Ammonis 3 (CA3) and Cornu Ammonis 1 (CA1) of the Hippocampus [15]. Changes of place fields after environmental modifications during experiments suggest that they are influenced by visually perceived geometrical information [16]–[18]. Furthermore, PCs were found to integrate non-visual afferents [19], [20]. Due to the stability of place fields over time, PCs are considered to memorize spatial locations [21]. In addition, they were found to be crucial for goal-directed navigation [22]–[26].

N. Waniek is with the Neuroscientific System Theory Group (NST), Technische Universität München, Arcisstraße 21, 80333 München, Germany. This work was partially supported by EU FET project GRIDMAP 600725.

PCs are subject to certain temporal processes and effects. Consolidation of spatial and episodic memories appears to happen during Sharp Waves and Ripples (SPW-R) [27]–[30]. These are events in which temporally compressed sequences of PCs are re-played in the order in which they were perceived. In addition, a recent study found pre-play activity of PCs in awake but stationary animals [31]. The study reported that sequences of PCs from the current location of an animal to a target location were pre-played in the order in which they were likely to be walked along afterwards, hinting to a path planning operation. Other studies reported PC activity in running animals which is temporally relative to Theta, an oscillation in the Local Field Potential (LFP) at 4 – 10 Hz [32], termed Theta Phase Precession (TPP) [33], [34]. Theta is believed to either provide or be the observable effect of synchronization of neural activity within and beyond the Hippocampus, thereby improving memory consolidation [35].Synchronization with extra-hippocampal areas such as the Pre-Frontal Cortex (PFC) is necessary, for instance, for decision processes and spatial navigation [36], [37]. During TPP, the PC which corresponds to the current location of the animal is active at the peak of Theta. In addition, several future and past cells spike in order of the spatial location of their place fields during the downward and upward slope, respectively, of the oscillation. Some authors suggest that TPP provides a mechanism for temporal buffering [38], [39].

GCs are stellate cells of the rodent Medial Entorhinal Cortex (mEC) and one synapse upstream of PCs [40]. After their initial discovery in 2005, they were since also discovered in other species, e.g. mice [41] and bats [42], [43]. Opposed to PCs, their spatial correlate is expressed as multiple fields of activity with respect to an environment [44], [45]. Curiously, the grid fields of a single GC arrange in a near-perfect hexagonal tesselation of the environment [40]. Grid fields can be characterized according to their relative phase, orientation, and size within an environment [45]. Commonly, the quality of grid fields is numerically assessed by the gridness score [46], a value which is determined by auto-correlating all responses of a GC within an environment and subsequently measuring how well peaks in the auto-correlation map are distributed hexagonally. It was discovered that grid field sizes increase in discrete steps along the dorsoventral axis, thereby forming several scales of representation [47]. Remarkably, the scales were reported to increase approximately by the factor 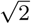 within and even across animals. Cells of the same scale are said to be part of one grid module, and the responses of one module densely cover the entire environment [40], [47].

Apparently, grid fields are influenced by the geometry of the surrounding environment [48]–[52], and a shearing effect was observed in regularly bounded environments [53]. Although GCs need excitatory afferents from the Hippocampus [54] and reportedly depend on Theta in the rodent mEC [55], they were also found to require converging inputs from visual pathways [56]. Furthermore, a somewhat overlooked statistical analysis reported that GCs tend to fire in correlation with an animal’s head direction and not with its movement direction [57]. The function of the hexagonal pattern in one and multiple scales is believed to provide a metric representation of space [44].

Conclusively, the HF is important for spatial information processing [11], [22], [58], [59]. In addition, it takes a critical role in the formation and retrieval of episodic memories [59]–[65]. It is structurally organized to form several loops of information processing, shows intricate local synaptic circuits of both excitatory and inhibitory neurons, and expresses significant amounts of recurrent connectivity [64], [66]–[68]. In particular, the inter-connectivity suggests that areas CA3 and CA1 form an auto-and hetero-associative memory, respectively [69]–[74]. While auto-association is suitable for storage and retrieval of patterns even when presented only with partial information, hetero-association allows storage of mappings from inputs to target states [75]–[77]. In combination, they can be used to store sequences of data [78].

### A. Overview of existing models and theoretical investigations

The origin of the hexagonal grid fields as well as their discretized field sizes are discussed controversially. Several models were developed to explain the peculiar arrangement and produce phenomenologically similar responses to real cells [79]–[81]. In Continuous Attractor Neural Network (CAN) models, the hexagonal fields appear due to temporal dynamics and recurrent lateral connectivity [82]–[84]. Recently, indirect evidence in favor of CAN models was presented by Yoon et al. [85]. In Oscillatory Interference (OI) models, the hexagonal patterns emerge due to integration of oscillatory afferents which depend on the Theta rhythm [86]. Although evidence suggests that Theta is required for the stable formation of GCss [87], the necessary accuracy of the oscillation in OI models was not reported so far. Other models use spatially modulated input, e.g. in form of PC activity, to drive a self-organizing process for the hexagonal arrangement [23], [88], [89]. They are supported by the observation that GCs require driving input from the Hippocampus [54].

Theoretical analyses found that Bayesian inference can be used to decode GC activity of multiple scales for the purpose of localization [90]. Although this encoding outperforms PCs [91], it is often assumed that the combination of several scales of GCs converge to form PCs [41], [92]. Another study demonstrated that a scale increment of 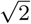 ideally covers a two dimensional input space when used for localization [93]. Other investigations used the hexagonal pattern for path integration [83], [94]. Furthermore, it was shown that the hexagonal encoding lattice can perform error correction during localization [95].

### B. Motivating questions

The theoretical investigations and existing models provoke several concerns. First, a redundant encoding for localization in both GCs and PC is unlikely due to energetically expensive maintenance of neural networks [96], especially when localization in GCs outperforms PCs [91]. Numerous evidence is in favor that PCs perform localization [3], [12],[22], [97], rendering the purpose of GCs under-determined. Second, the hexagonal encoding yields ambiguities when used as a path integration mechanism, and requires stabilization to prevent or reduce drift due to noise in real-world scenarios [98]. Other integration schemes such as a homing vector or a pedometer appear more likely. Third, only few GCs models address temporal aspects of spatial information [99]. However, the Hippocampus is known to form a basis for episodic memory [60], [62], [63], [100], and GCs appear to require episodic data [101]–[103].

The goal of this work is to address these concerns. Given that the Hippocampus and Entorhinal Cortex (EC) are required for goal-directed navigation [104], [105], it is proposed that GCs efficiently encode spatial transitions in sequences of locations.

### C. Organization and brief summary

The paper is organized as follows. First, a model for self-organizing GCs is presented in which each GC learns as many valid transitions as possible via dendritic computations. The model is evaluated in simulations of competitive, recurrently coupled GCs. Grid fields emerge quickly and remain stable over long simulation times.

Next, transition encoding is examined in the light of computational performance and behavioral relevance. Here, a novel scale-space model of GCs is proposed to achieve behaviorally significant run-times. In particular, it is first shown theoretically that the optimal increment across scales is 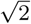. Then, it is demonstrated that multiple scales improve runtimes exponentially in a simplified model. To form the scale-space representation, temporal buffering of spatial locations is required.

Subsequently, the formal theory for transition encoding on which the self-organizing model is based is presented. Here, goal-directed navigation is examined from a logical perspective in which sequences of locations and transitions are formalized mathematically. Several constraints are introduced which are necessary to form consistent sequences. To logically examine transition encoding, inspiration was taken from the analysis of time in distributed computing systems [106], and from approaches to understand and model causality in theoretical computer science [107]. Furthermore, a notation for transitions is used which is based on Communicating Sequential Processes (CSP) [108]. The block concludes with proving that a hexagonal arrangement of transition encoders is optimal for two dimensional Euclidean space with tools and methods from graph theory.

Finally, the results are discussed in detail. They are examined with respect to inlfluential related work and biological findings, testable predictions are made, and links to other areas of research are established.

## II. SELF-ORGANIZING GRID CELLS FOR TRANSITION ENCODING

A formal theory for sequences and transition encoding is introduced in Section IV. Briefly summarized, the results are as follows. The theory defines transition bundles which encode as many transitions from one symbol to another. Thereby symbols and bundles can be used to generate sequences, for instance for goal-directed navigation. To generate valid sequences, a bundle cannot associate to starting points if it already encodes a transition leading to this point. Given the assumption of a suitable sensory input, it is shown that the optimal distribution of bundles to minimize their number is hexagonal in Euclidean space, and that the minimal number is three.

The results of the formal theory are now used to derive a biologically plausible model of self-organizing GCs. Each cell corresponds to a transition bundle that encodes transitions between spatial locations. The necessary behavior of each cell, i.e. associating with as many presynaptic states as possible, is modeled by dendritic computation in form of multiple dendritic spines, or short *dendrites*.

However, a single cell must decorrelate from locations to which one of its memorized transitions leads. In Euclidean space, the target region of a transition is a volume around the current location. Conceptually this is modeled as receptive field dynamics with on-center and off-surround regions, the first corresponding to the start location of a transition and the latter to the target region. The on-center off-surround receptive field is expressed by each dendrite individually. Figure 1a illustrates the dendritic tree of a single cell as well as the preference of cells to correlate with multiple inputs in two dimensions.

**Figure 1:**
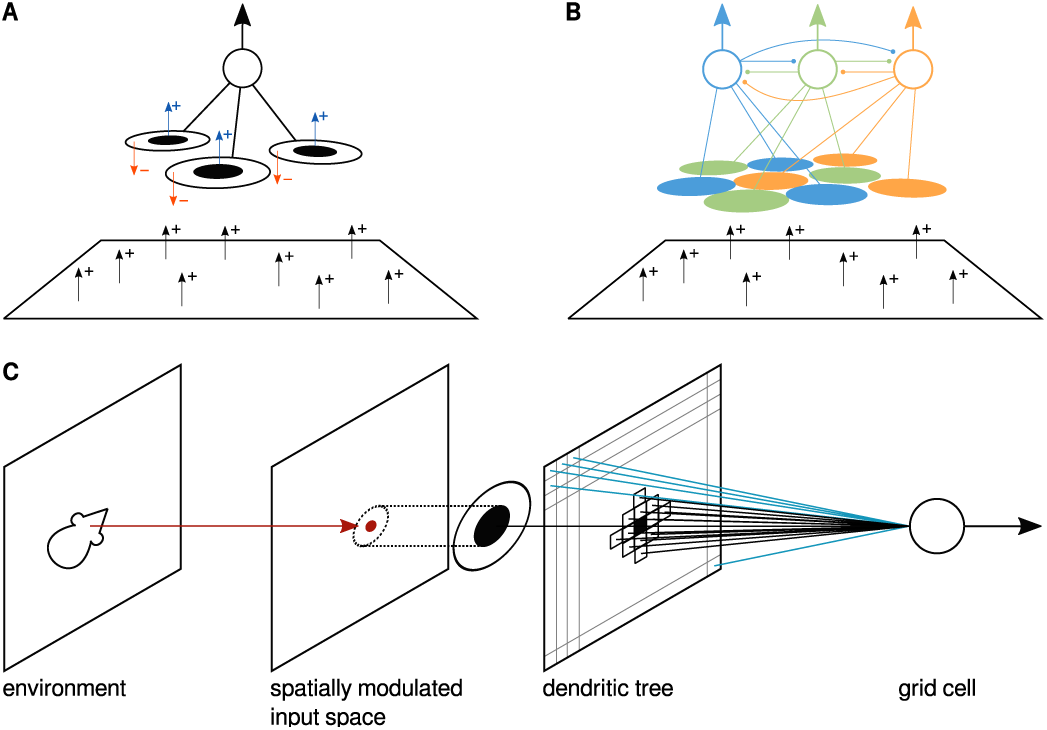
Conceptual overview of the model and detailed view of the dendritic organization. (a) A single cell shows preference to associate with as many inputs as possible, indicated by black arrows with a + sign. Transition constraints are expressed as dendritic computation in form of on-center off-surround fields, indicated by small circles and arrows with – and + signs. (b) To cover an entire input space, at least three cells are required which organize their fields due to competitive dynamics. (c) The location of the animal induces a singular response (red circle) in a spatially modulated input space. Specifically, coordinates are used to during simulations. The dendritic tree of each grid cell covers the entire input space, and dendrites are organized on a regular lattice according to the image (indicated by little boxes and gray lines). Each dendrite expresses a receptive field of given certain size (circles with black center), which may overlap with neighboring dendrites. Therefore, multiple dendrites may get activated (black small boxes) for a single stimulus. All other dendrites remain silent (blue lines).

Due to the decorrelation, a single cell is unable to capture transitions from arbitrary locations in the environment. Any transition which starts at a location from which the cell decorrelated has to be covered by another cell. In concordance with the theoretical results presented in Section IV, the system is modeled as a minimal network of *N*_*g*_ = 3 cells, illustrated in Figure 1b. To preserve uniqueness of transitions and their corresponding starting locations, the interactions between cells have to be competitive.

### A. Assumptions and model specification

Each GC samples an input space with *N*_*d*_ dendrites. Interactions between a GC and presynaptic afferents are modeled as follows, based on several assumption. For instance, it is assumed that any location in the experimental environment induces a unique sensory representation. A likely candidate for this purpose is the Boundary Vector (BV) space [18]. However, coordinates are used to reduce the complexity of the model.

The location *x* of the animal is thought to evoke spikes from several presynaptic neurons with overlapping tuning curves. The spikes are integrated by dendrites *i* which encode locations not further apart than σ_1_, i.e. |*x*_*i*_ *-x*| ≤ σ_1_. The totality of these dendrites, denoted as the set *B*_*n,t*_ for cell *n* at simulation time *t*, is assumed to be sufficient to drive the GC to its spiking threshold. In addition, dendrites *j* which encode nearby locations are expected to generate an excitatory post-synaptic potential shortly after the GC fired, i.e. σ_1_ *|x*_*j*_ *-x|* ≤ σ_2_ and denoted *C*_*n,t*_. In the simulations, the temporal integration of presynaptic spikes relative to the spike time of a GC is collapsed into one singular time-step. The simplification can be understood as a binarization of Spike-Timing Dependent Plasticity (STDP) dynamics. Consequently, the set of all dendrites which receive stimulation at time step *t* for cell *n* is the set *D*_*n,t*_ = *B*_*n,t*_ *ᵁ C*_*n,t*_, and corresponds to the formation of an on-center and off-surround receptive field. The on-center represents the location at which a transition starts, and the off-center region the target area to which a transition leads.

Due to the correspondence of the input space with coordinates, each dendrite *i* can be identified by a coordinate *x*_*i*_.The dendrites are organized on a regular lattice, and each dendrite *i* of cell *n* stores a weight *w*_*n,i*_ ϵ [0, 1] that indicates the dendrite’s probability to associate with the corresponding input. The entire vector of weights is denoted as *w*_*n*_. For the simulations presented below, the dendritic tree reduces to a two dimensional sheet of weights which covers the entire input space. The processing of presynaptic afferents, dendritic computation, and dendritic organization of a single cell are illustrated in Figure 1c.

Activation *a*_*n*_ of a single cell *n* is computed according to Equation 1 as the weighted sum over all dendrites *B*_*n,t*_ which receive stimulation at time-step *t*. Weighting only appears on the dendritic side, i.e. presynaptic activity is considered to be binary.

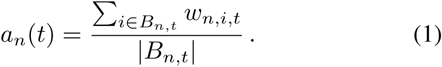

Each cell maximizes the number of locations to which it is associated. This is formalized as a function *L*_*n*_(*w*_*n*_), specified in Equation 2, which tells the *dendritic load* of a cell. It can be understood as a diffusive *eagerness* of each dendrite to associate with arbitrary presynaptic inputs.

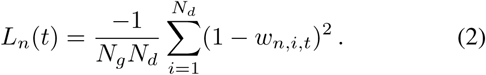

Furthermore, a network of *N*_*g*_ cells has to minimize co-activation *K*, modeled according to Equation 3.

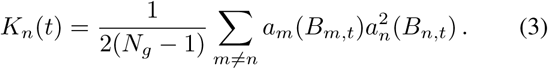

In addition to diffusive dendritic load, each cell has to express specificity for transitions. A cell which associates with a certain presynaptic state has to dissociate from nearby or similar states. Clearly this information is present in the on-center and off-surround dynamics of the dendritic computation described above. The contributed errors for the on-and off-portions are denoted *E*^+^(*t*) and *E*^*-1*^(*t*), respectively. Intuitively, the error terms can be understood as follows. Consider the term *E*^-^(*t*).Any dendrite *i* which synapses with erroneous presynaptic input, i.e. a state to which a transition that is encoded by the cell leads to, contributes an error proportional to *ω_n,i,t_*. The errors are minimized according to the derivatives specified in Equations 4 and 5, respectively.

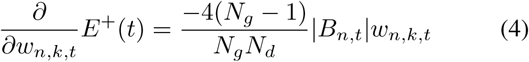

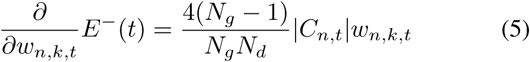

The derivatives are inspired by the instar learning rule [109]. Hereby, minimization depends only on *ω_n,k,t_* and the total amount of stimulated dendrites *|B*_*n,t*_*|* and *|C*_*n,t*_*|*. Specific crosstalk with potentially remote dendrites *ω_n,i,t_, i ≠ k*, perceived to be biologically unlikely, is thereby avoided.

Summarized, a GC is characterized by the error function *F*_*n*_ of Equation 6.

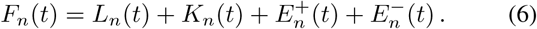

The error function can be derived with respect to a weight *ω_n,k,t_* to determine a gradient descent learning rule to minimize the error over time, shown in Equation 7.

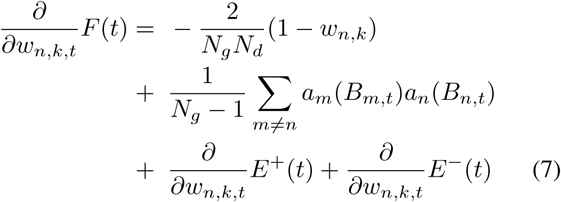

Changes to dendritic weights according to Equation 7 are subject to global winner-take-all and local activation mechanisms. All cells perform weight updates with respect to *L*_*n*_(*t*) and *K*_*n*_(*t*), however limited to dendrites *D*_*t*_. Furthermore, only the most active GC, i.e. if *n* = arg max_*m*_ *a*_*m*_(*t*), receives nonzero gradients *E*^+^ and *E*^-^. The gradients *E*^+^ and *E*^-^ are only applied to corresponding dendritic weights, i.e. only dendrites of *B*_*m,t*_ receive a gradient for *E*^+^ and only dendrites of *C*_*m,t*_ a gradient for *E*^-^. Thereby, changes are local to dendrites which perceived presynaptic activity and proportional to their individual weights. Subsequently, the update of a weight *ω_n,k,t_* is modulated non-linearily. Finally, all weights are rectified, i.e. weights are clamped from below to zero, indicated by the bracket notation [·]^+^. Conclusively, a weight update follows according to Equations 8, 9 and 10.

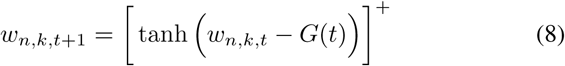

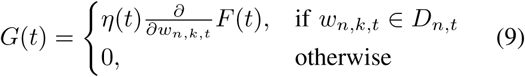

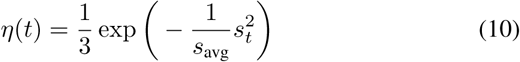

The learning rate (*ηt*) depends on the current speed *s*_*t*_ in 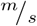 of the animal as well as the average speed *s*_avg_. This dependence models increased uncertainty of perception. It is assumed that an animal which moves at a high speed is less likely to identify locations perfectly. Correspondingly, the accuracy of learning transitions declines with increased speed, expressed by a lowered learning rate. In the simulations presented below, the average speed *s*_avg_ was extracted from precomputed trajectories. However, a moving average is expected to yield qualitatively similar results.

### B. Simulation setup and results

A dendritic weight is initialized to 1.0 with a probability of 0.1, or set to zero otherwise. Input to the model was presented in form of the speed of the simulated animal to adjust (*ηt*), as well as the location to drive the presynaptic state. The simulated animal moved throughout the environment with movement statistics close to real recordings, extracted from recordings by Hafting et al. [40]. An example of a simulated trajectory as well as statistics of the distribution of speeds and angular velocities are given in Figure 2. The trajectories were pre-computed to extract *s*_avg_. In each simulation, the animal started in the middle of the environment. The network model was simulated 400 times for the minimal number of *N*_*g*_ = 3 cells per network, each cell exhibiting *N*_*d*_ = 48 dendritic weights. Dendritic stimulation due to presynaptic activation was governed by the parameters σ_1_ = 0.10, and σ_2_ = 2σ_1_.

**Figure 2:**
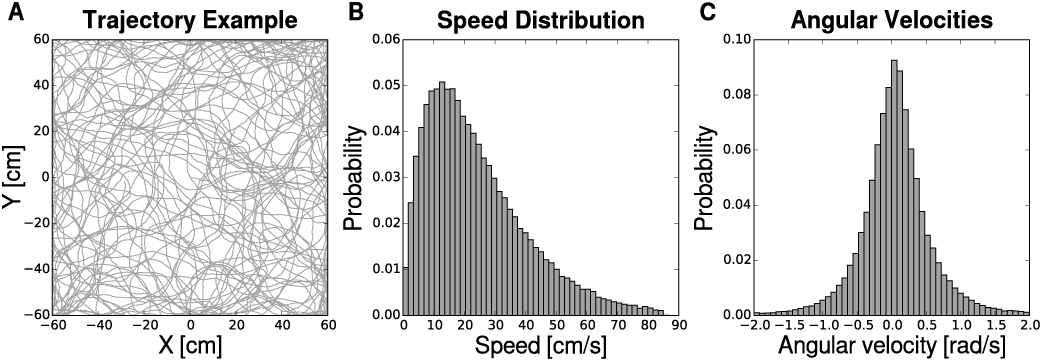
Realistic trajectories (a) were generated by tuning the distribution of speeds (b) as well as angular velocities (c) to resemble real recordings.

The gridness score of a cell was calculated without generating spikes. The value was extracted directly from the dendritic weights. All weights *ω*_*n,k*_ ϵ [0, 1], thus they encode a probability to spike given a certain input. Consequently, the distribution of spikes precisely follows the dendritic weights. The gridness score was computed according to Sargolini et al. [46], however without previously smoothing the weight maps. Furthermore, the orientation of the dendritic weights was computed following the methods described by Hafting et al. [40]. The relative grid orientation error of the network was computed by accumulating the mutual differences of orientations between all cells.

Both, gridness scores and relative orientation errors, are presented in Figure 3 for 400 simulations with a total simulated duration of 180 minutes. The top row of the figure shows that cells with a gridness score above zero appeared early in most of the simulations. Furthermore, the bottom row demonstrates that relative grid orientation errors declined over time.

**Figure 3:**
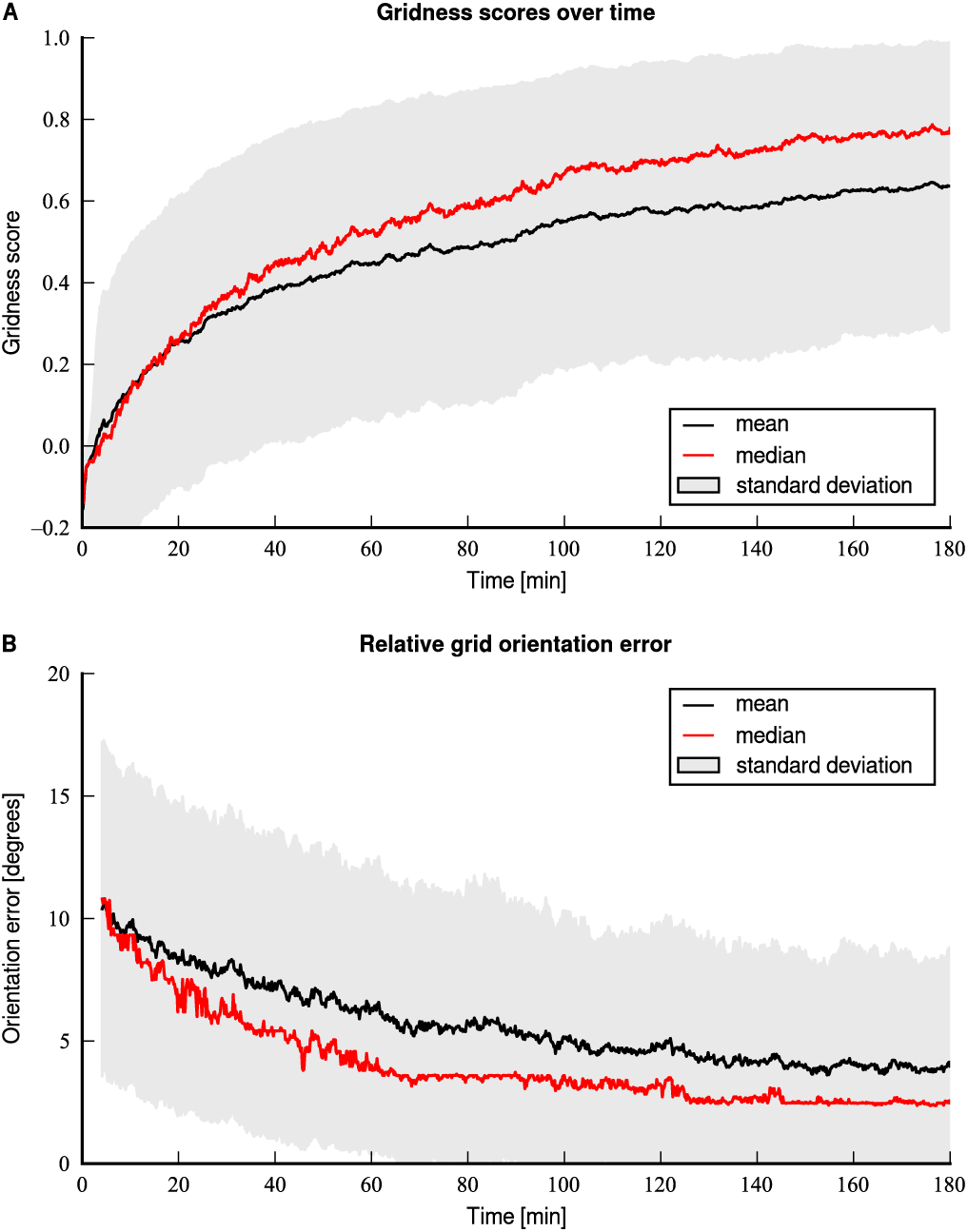
Gridness scores and orientation over time for 400 simulations. (a) Gridness increases over time, the median stays above zero already after approximately 2.5 minutes of simulated time. The mean remains positive after about 4 minutes. (b) Relative grid orientation errors decline over time. The first four minutes of data were cut off to allow the cells to form detectable peaks in their weight distributions.

Examples of three simulations, one with a high final average gridness score, one with mean score, and one with low score, are depicted in Figure 4. The figure displays the dendritic weights of each of the neurons at several points in time during the simulation in form of heat maps. The first column shows the dendritic weights after a few iterations of the system and not immediately after initialization, visible in form of slightly increased values in the central weights, and that the weight maps are presented as-is and were not post-processed with a smoothing filter. Each panel contains a small inlay displaying the gridness score of the presented weight distribution. The emerging weights are non-binary and smooth despite the binary presynaptic activation of the dendritic tree. In addition, the weights at time *t* = 180 min indicate effects similar to recently reported distortions such as skewing, or misalignments with walls [53]. Analysis of these effects is left for future work, though. The last column of Figure 4 shows the auto-correlogram of the dendritic weights at the end of each simulation. The detected orientation of the arrangement of weights is displayed in form of a black bar from the central peak to the first peak above the horizon in counter-clockwise direction.

**Figure 4:**
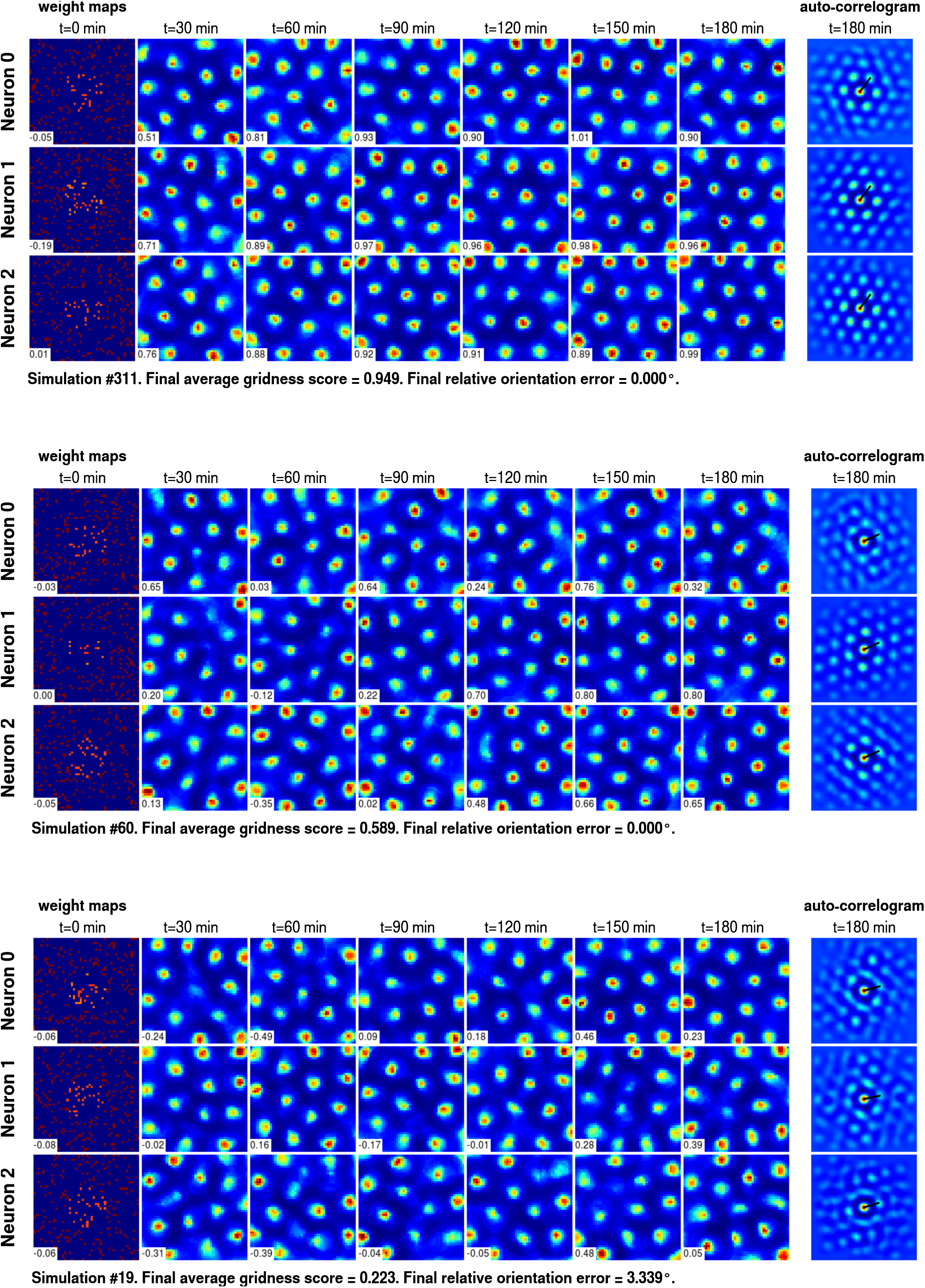
Examples of grid cells at several points in time during the simulation and final auto-correlograms with indicated grid orientation. White inlays with numerical values show gridness score of the respective weight map.

## III. A SCALE-SPACE MODEL OF GRID CELLS

The entorhinal-hippocampal loop is proposed to form an Multi-Transition System (MTS) for both episodic sequences as well as goal-directed navigation, and stores symbols and transition bundles as follows. A neuron representing a spatial symbol can be understood as a PC, hence the terms symbol and PC are used interchangeably. Novel symbols are acquired in a neural associative memory *M*_Σ_, episodic and actually performed transitions are recorded as bundles in *M*_Π_, while purely spatial bundles are subject of *M*_Г_. Spatially modulated afferents, required for the recruitment of new symbols and transitions, are represented by neural memory *M*_Δ_. The interactions between these memories are depicted in Figure 5d and focus of the remainder of this section. The model is inspired by previous work by Cuperlier et al. [110], [111] and by Hirel et al. [112].

**Figure 5:**
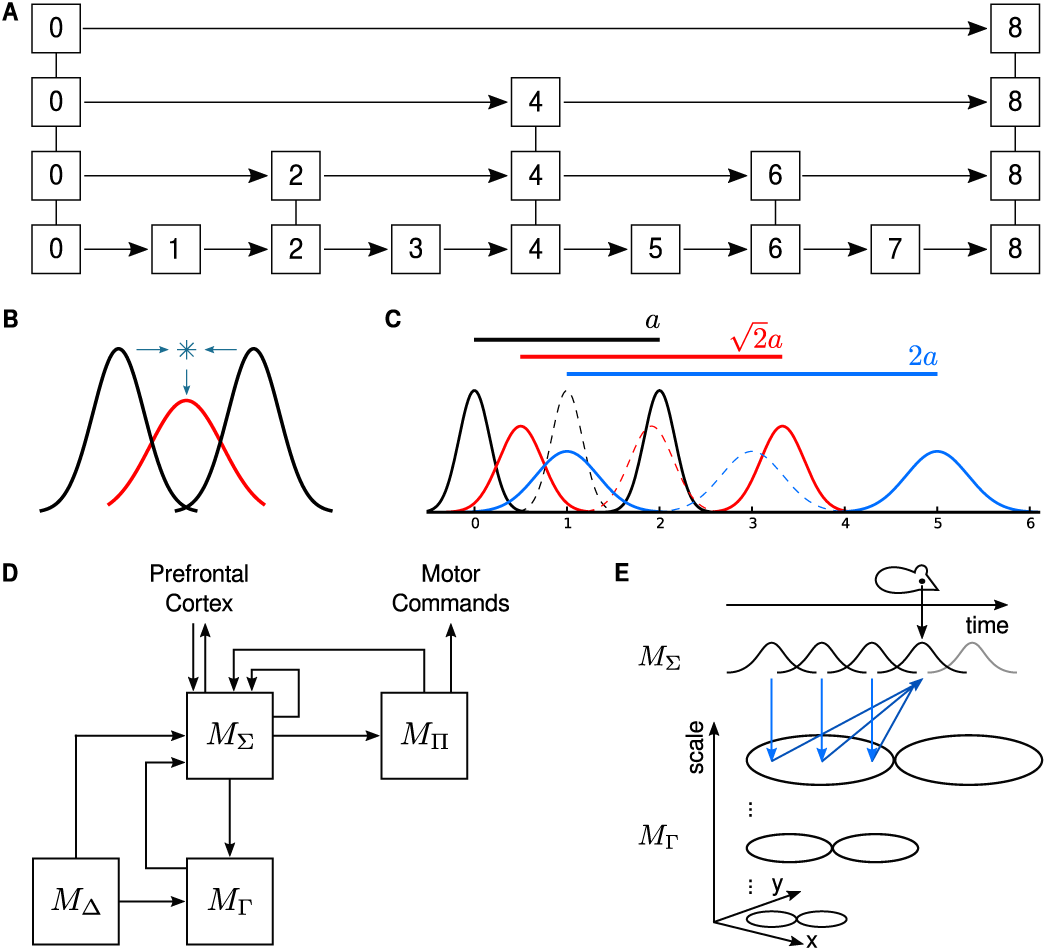
Skip list, scale-space model construction, overview, and learning. (a) A skip list for discrete data is a data structure which uses a hierarchy of *fast lanes* to accelerate search [114]. In the ideal case illustrated here, the hierarchy guarantees logarithmic time, i.e. exponential speed up of retrieval, and corresponds to search in a binary tree. For fast lane traversal, comparison of items is required. (b) For probabilistic data represented in a neural network, convolution of two consecutive normally distributed probability density functions (pdf) is required. This yields another pdf with variance increased by factor *√*2. (c) Recursively applied this constructs a scale-space representation. The scale space allows to compare items across larger distances of the represented space, similar to the comparison on fast lanes in a discrete skip list. Dashed lines indicate off-surround areas. (d) The final model consists of the four memories *M*_Δ_, providing spatially unique information, *M*_Ʃ_, storing spatial symbols or place cells, *M*_Г_, which learns spatial transitions between perceived locations, and *M*_ᴨ_, storing actually performed or *temporal* transitions between symbols. The memories *M*_Ʃ_ and *M*_Г_ directly correspond to the items and transitions of a discrete skip list. The model was inspired by previous work by Cuperlier et al. [110], [111]. (e) During learning, acquisition of large-scale transitions requires to access locations and symbols which were perceived in the past to learn the association with the currently active symbol. Input from *M*_Δ_, which is co-active with *M*_Ʃ_ during learning, is omitted in the figure for clarity, and that only the on-center regions of transition encoders of *M*_Г_ are shown for the same reason.

The system can be queried to expand the path *σ*_*s*_ *⇝* σ_*t*_ by recursion. After activation of a start symbol *σ*_*s*_ in *M*_Ʃ_ all corresponding transitions activate in *M*_ᴨ_ and *M*_Г_. Recursively this leads to activation of subsequent symbols in *M*_Ʃ_ due to the bi-directional connection between *M*_Ʃ_ and the two transition memories. Simultaneous co-activation of multiple symbols or transitions is allowed. Hereby the system is able to expand *σ*_*s*_ ⇝ *σ*_*t*_ into known paths in parallel until the target state *σ*_*t*_ is activated in *M*_Ʃ_, or until a maximal number of recursions is reached.

### A. Multi-scale transitions and the number 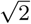

Both the outlined model above as well as the ones published by Cuperlier et al. [110], [111] or Hirel et al. [112] have a biologically significant issue. Consider the computational and thus behavioral implications of the recursive retrieval in the transition system for an animal which has to travel 200 m in a straight line from a feeding site back to its home location. The animal is equipped with an entorhinal-hippocampal transition system for path planning which acquires novel spatial symbols every 20 cm. Thus, 1000 spatial symbols cover the path from start to goal, ignoring overlapping place fields. Furthermore, assume that temporal dynamics and delays for spike generation and propagation consume 10 ms of time per memory. After activation of a symbol in *M*_Σ_, recursive retrieval with the help of *M*_ᴨ_ and *M*_Ʃ_ therefore requires 20 ms of time to activate a subsequent symbol in *M*_Σ_. In total, this accumulates to 20 s of time to query the existence of an expansion – if it exists – in which the animal may fall victim to a roaming predator. Clearly an acceleration technique is required.

From the perspective of computer science, symbols and transitions stored in a transition system form an ordered list of data. Given random access to elements, binary search is known to demonstrate the asymptotically optimal runtime [113]. For lists without random access, skip lists were developed [114]. The data structure builds a hierarchy of look-ahead links, often called *fast lanes*, by which search is accelerated exponentially. Given uniformly sampled elements of data, skip lists express an immediate duality to binary search [115]. In particular, the hierarchically structured fast-lanes exponentially decrease the search time for a range query on discrete and equidistant data. The look-ahead distance is governed by a factor of two, depicted in Figure 5a. Two consecutive locations have to be compared and combined as fast lane to form the next level of transitions in this hierarchical approach.

How does this relate to the entorhinal-hippocampal loop? The value represented or detected by a single real neuron depends on its tuning curve, which is usually bell-shaped [116], [117]. Furthermore, the perception of two consecutive locations is assumed to depend on optimally sampling an input space. Due to a finite number of neurons to represent sensory stimuli, it can be assumed that there is a minimal resolution to discriminate between two (sensorily) adjacent positions, called *eigenresolution* and denoted *σ*_eigen_ and σeigen in the one and multidimensional case, respectively. In other words, two locations can be distinguished if they are at least *σ*_eigen_ apart in the sensory space. For ease of comprehension, *σ*_eigen_ can be understood to correspond to a distance in Euclidean metric space, similar to the parameter *σ*_1_ used in Section II. Conclusively, neural afferents to GCs are proposed to be described by an *n*-dimensionally normally distributed Probability Density Function (pdf) according to Equation 11.

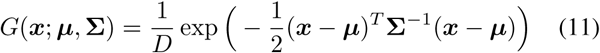

The pdf is parametrized by the preferred stimulus *µ* and tuning width expressed by the covariance matrix Ʃ. The normalizer *D* is specified according to *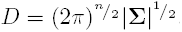.*

Combining two locations to construct a transition lookahead requires to convolve two consecutive spatial samples, parametrized by *µ*_1_, *µ*_2_, and denoted as Σ_1_, and Σ_2_. Combining distant locations would violate the coherency constraint of Multi-Transition Theory (MTT), presented in Section IV. It can be shown that convolving two normally distributed pdfs results in yet another normally distributed pdf with combined mean and co-variances (see supplementary material to Vinga et al. 118] for details). In particular, the resulting pdf is parametrized according to *µ* = *µ*_1_ + *µ*_2_ and ∑ = ∑_1_ + ∑_2_. Given uniform input sampling and symmetry, i.e. ∑_1_ = ∑_2_ = ∑ eigen and ∑_eigen_ = *σ*_eigen_***I***, it follows that the variance *σ*^2^ of a sampling process for spatial look-ahead is *σ*^2^ = 2σ_eigen_^2^. Thereby the integration area of a spatial sampling process *doubles* during the construction of an additional scale to perform transition look-ahead, regardless of the dimensionality of the pdf. The convolution process and three consecutive scales are depicted for the one-dimensional case in Figure 5b and c, respectively.

Conclusively, the radius of the spatial sampling process ideally increases by a factor of 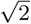 from the first scale to the next. In concordance, the spatial period of transition encoders which perform spatial sampling as part of their dendritic computation increases by 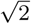. Repeatedly applied, the technique forms an entire stack of probabilistic look-ahead transition encoders with a discrete scale increment by a factor of 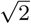. This is similar to the *fast lanes* of skip lists in the asymptotically optimal case, however with the difference of using probabilistic instead of discrete data.

### B. Implementation details of a proof-of-principle model

The theoretical results were simulated in the following deterministic manner as proof-of-principle. The model consists of the four memories *M*_Δ_ to represent input, *M*_Σ_ to store symbols, *M*_Π_ to record temporal transitions, and *M*_Г_ for the storage of spatial transitions, interacting according to Figure 5d. Each memory sacrifices biological accuracy to focus on the effect and requirements of using and learning a scale-space model of transition encoding.

Likewise the input to the model of Section II, the memory *M*_Δ_ reports the coordinate of the animal in two dimensional space to reduce the complexity of the model. The memories *M*_Ʃ_ and *M*_ᴨ_ are implemented as artificial neural networks similar to growing neural gas [119]. *M*_Σ_ associates with presynaptic states of *M*_Δ_ and thereby corresponds to PCs, whereas *M*_Π_ records actually performed transitions from one neuron of *M*_Σ_ to another. Thus, any neuron in *M*_Π_ associates presynaptically with a single symbol neuron of *M*_Σ_, and recurrently connects back to all symbols in *M*_Σ_ to which transitions were detected. A novel transition neuron is recruited in *M*_Π_ only when the corresponding symbol of *M*_Σ_ is not yet known to *M*_ᴨ_. Recruitment of novel neurons in *M*_Σ_ is determined by the difference between the afferent from *M*_Δ_ and the value represented by the best matching unit. Given the use of Euclidean coordinates in *M*_Δ_, this is simply the distance between the respective coordinates. In the results presented below, recruitment of novel PCs was triggered when the best matching unit represented a value that was displaced by at least 10 cm from the current location of the animal.

The distribution of optimal transition bundles in memory *M*_Г_ is predetermined to simplify the simulation. Instead of learning the optimal distribution of the bundles similar to Section II, hexagonally distributed Voronoi cells with a radius of 20 cm are used on top of the coordinate space provided by *M*_Δ_, and the scale-increment from one scale to the next is determined by the factor 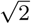. Although absolute areas in form of Voronoi cells neglect the probabilistic nature of neural encodings, used to derive the optimal scale increment of 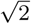, it is believed to suffice for demonstration of proof-of-principle.

Learning look-ahead transitions requires to buffer symbols temporally. Identification of locations and recruitment of symbols in *M*_Σ_ depends on afferents from *M*_Δ_, similar to the smallest scale of GCs in *M*_Г_. Without additional information, a symbol is thus only active in a place field that is expected to depend in size on the resolution *σ*_eigen_ of the sensor representation. However, any PC which falls into the on-center region of a large-scale GC indicates the start of a look-ahead transition. Similarly, each PC which is active immediately after the large-scale GC has to be considered a target of a look-ahead transition. Consequently, a large-scale GC has to associate with several previously experienced PCs before spiking and dissociate from prospective PCs to learn look-ahead transitions. Hence, all symbols of *M*_Σ_ are buffered in consecutive order to allow learning, illustrated in Figure 5e. Conclusively, learning of transitions in *M*_Г_ is gated by co-activation of afferents from *M*_Δ_ and temporally buffered information from *M*_Σ_. The biological feasibility of buffering as well as co-activation learning will be discussed further below.

The sensor representation *M*_Δ_ is not required for retrieval, only the memories *M*_Ʃ_, *M*_ᴨ_, and *M*_Г_ are recursively iterated. While co-activation of *M*_Σ_ and *M*_Δ_ during look-ahead learning is implemented as a *logical and* operation, retrieval requires to toggle this modality to *logical or* to allow *M*_Σ_ to drive *M*_Г_ by itself. Recursion is performed until the neuron which corresponds to the target symbol is activated in *M*_Σ_, or until a maximal number of recursive invocations is reached. Furthermore, querying the existence of a trajectory performs retrieval across all scales concurrently. For instance, assume that a transition between two locations was learned in the largest scale. Certainly this means that a trajectory is known which links the two locations on the smallest scale across several intermediate places. However, these small-scale details of the trajectory are not necessary to merely determine the existence of a viable path, or can be retrieved while the animal is already walking in the general direction of the goal. Thus, a large scale transition encoder activates all corresponding distal symbols concurrently to any other symbols that are activated due to smaller scales of transitions.

### C. Simulation results

The deterministic model was evaluated on an S-shaped trajectory in a square environment, depicted in Figure 6a. To observe the impact of temporal buffering, two slightly different versions of the model were examined. In the first variant, temporal buffering was ignored and random access to all symbols was granted during learning. For instance, learning a feasible spatial transition only depended on the spatial vicinity between locations. The second variant required that previous locations were in the temporal buffer in a suitable time window. The S-shaped trajectory was designed such that any effects of the temporal buffer are recognizable when using at least one additional transition scale, illustrated in Figure 6a-c.

**Figure 6:**
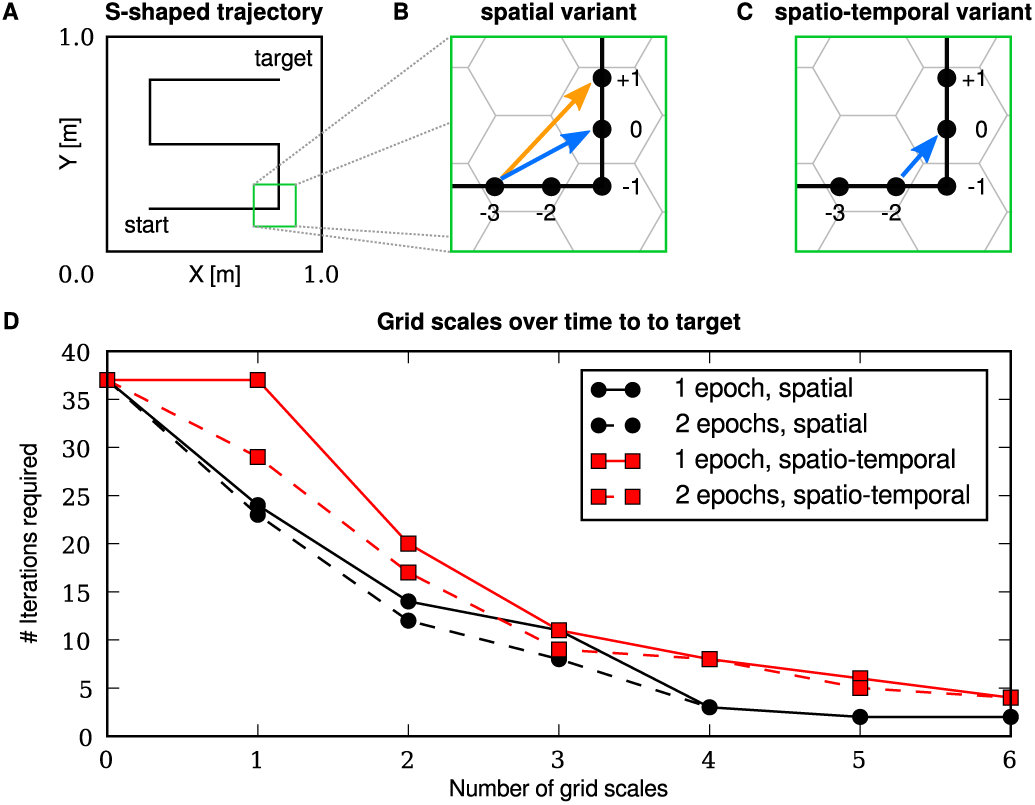
Expected behavior and results of the scale-space model. (a) The S-shaped trajectory which was used during evaluation. (b + c) The zoom in on one of the corners of the trajectory shows the difference in behavior between the spatial and spatio-temporal variant when one additional transition scale is used. Black dots indicate place field centers, the numerical value closest to each dot the relative time of perception with negative values going to the past. In the spatial variant, transition learning is only gated by spatial vicinity. During the initial exploration, a transition is thus learned from place cell at time –3 to 0 (blue arrow). During the second exploration, a transition from –3 to +1 is learned due to prefetching (orange arrow). In the spatio-temporal variant, learning is restricted to items in a suitable temporal buffer. Consequently, only the transition from the place at –2 to 0 is learned. The hexagonal Voronoi cells only depict the on-center of transition encoders. (d) Using multiple scales of grid cells improves the run-time, measured in number of iterations, exponentially to determine if a viable path from start to target exists.

Each variant was evaluated after only one trial during which learning was disabled. Furthermore, each variant was examined in a second trial with learning enabled during which the temporal buffer pre-fetched future symbols from *M*_Ʃ_ that were acquired during the first trial. After the second trial, learning was disabled again and replay of the trajectory was recorded.

Results are depicted in Figure 6d, which shows the number of recursive iterations required until the target location was found for increasing numbers of additional scales. The results show that the number of necessary iterations declines exponentially with an increased number of scales, thus improving the runtime of the model. In addition, the impact of temporal buffering is discernible, i.e. the spatio-temporal model performs slightly worse than the purely spatial model. However, this is considered to be an artifact of the simplified model of GCs that was used. Furthermore, mental travel during the second exploration of the environment improves the runtime of the model. The reason is prefetching of future locations in the temporal buffer structure combined with learning.

## IV. A FORMAL THEORY FOR MULTI-TRANSITION SYSTEMS

Spatial navigation can be viewed as a process in which sequences of symbols are learned and produced. For this purpose, knowledge about feasible transitions from one symbol to another is required. Goal-directed navigation is therefore examined using the following axiomatic system, called Multi-Transition Theory (MTT), in which spatial locations are represented by symbols and movement from one location to another by transitions.

### A. Symbols, alphabets, and sequences

Consider an animal which moves through three rooms. The trajectory of the animal can be described the sequence of symbols *A, B, C*, e.g. each representing one room. The meaning of a symbol is not pre-determined, for instance the symbols could also represent the *event of perception* of each corresponding room. The entirety of symbols forms an alphabet and their consecutive ordering a sequence, both of which are captured in the following definition.

**Definition 1** (Alphabet and sequence). *An* alphabet Σ *is a finite set of symbols. A* sequence *(or word) is an ordered tuple of symbols σ*_*i*_ *ϵ Σ, i.e.* (*σ*_0_, *σ*_1_,…) = *w ϵ Σ* ^+^, *where*^+^ *is the Kleene plus operator.*

A trajectory of an animal is thus viewed as a sequence, and moves along several symbols. However, repetition of a single symbol is disallowed.

**Axiom 1** (Non-stationarity). *A sequence is non-stationary if any two successive symbols σ*_*i*_ *and σ*_*i*+1_ *are distinguishable,i.e.;σ*_*i*_ *≠ σ*_*i*+1_.

This does not limit general capabilities. Two consecutive but distinct symbols of a sequence can have the same associated meaning, for instance the perception of a certain room.

The directional ordering of a sequence is expressed using the arrow notation *→* For example, *A→B* means that the symbol *B* causally follows after symbol *A*. However, *time* is not immediately given in the definition and needs to be stated explicitly. Thus, symbols and transitions can be viewed as terms of propositional logic. For instance, *A → B* means that if *A* is *true*, it follows that *B* is also true. Thereby they form a *chain of causality*. In addition to *→* the arrow ⇝ exists, e.g. *A*⇝C means that there exists a *path* from *A* to *C* which bridges *n ≥* 0 intermediate symbols. The negations of the notation are *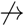* and *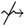*.

**Axiom 2** (Coherency). *Let ω* =*σ*_*i*_, *i ϵ* {0,…, *N*} *be a sequence of N symbols. ω is* coherent *if and only if σ*_*i*_ *→* 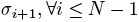.

Coherency is required in goal-direction navigation. Consider an animal which travels from a starting point to a target location. The animal has to construct a trajectory without any gaps to reach its destination. Otherwise, it may express undefined behavior or displacement activity as it does not know how to proceed to its goal. Without a distinct goal, the animal performs explorative movement in which novel symbols are acquired.

**Axiom 3** (Validity). *A sequence ω is* valid *or* acceptable *if it is both non-stationary and coherent.*

Using these notations and axioms, goal-directed navigation from a start *A* to a goal *C* can be expressed as a *program* which expands the path 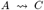 into any valid sequence *A → σ*_1_ *→… → σσ*_*N*–1_ *→ C*, if it exists. The next section will discuss how the expansion can be denoted and performed for arbitrary symbols

### B. On Universal Multi-Transition Systems

The arrow notation specifies relations between symbols. Consider the example *A → B* which contains the transition from *A to B* The transition is a tuple (*A, B*) which maps one symbol to another and known, for instance, from Reinforcement Learning (RL). There it is denoted as a transition function mapping from a set of states and a set of actions *R* to the next state, i.e: *τ*: ∑ ×R→∑[120]. The notation is now extended for the purpose of encoding multiple feasible transitions.

**Definition 2** (Transition system, set, bundle, and point). *A Multi-Transition System (MTS) ℳ is the pair*

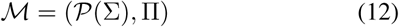

*where P*(∑) *is the power set of* ∑. *A set* Ω *ϵ P*(∑) *is called a* configuration *of ℳ. All symbols σσ*_*i*_ *ϵ Ω are considered to be* true.

*The set* II *is called* transition set *and contains sets π*_*i*_, *called* transition bundles. *In turn, a transition bundle π*_*i*_ *is a set of transitions 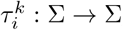called k*-th transition point of *π*_*i*_.

An MTS *ℳ* can be interpreted as a state machine in which multiple states can be active simultaneously. Indices will be dropped if they are clear from context. Figure 7a illustrates a sequence, transition points, and bundles. The following additional terminology and notation will be used.

**Figure 7:**
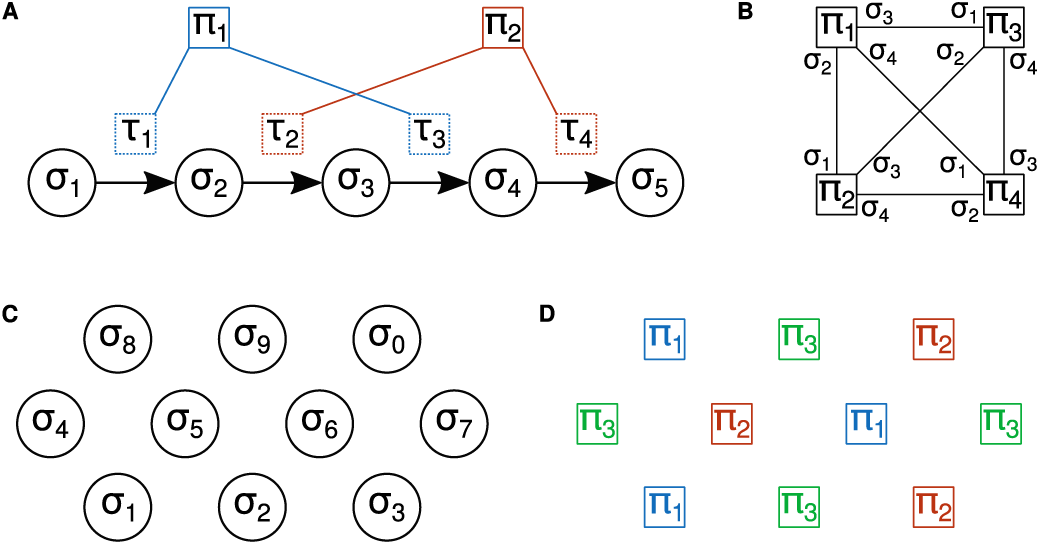
Transition graph examples. (a) A sequence of five consecutive symbols *σ*_1_,…, *σ*_2_. The transitions *τ*_1_,…, *τ*_4_ can be bundled into *π*_1_ and *π*_2_ without violating any constraints.(b) In a universal transition system, the transition graph is fully connected. By construction, each edge is associated with two symbols. (c) In the two dimensional case, the optimal distribution of symbols is a hexagonal arrangement [122].(d) The optimal distribution of transition bundles in the two dimensional case follows the distribution of the symbols. However, bundles can be repeated periodically in a hexagonal fashion. Edges in the graphs of (c) and (d) were omitted to improve clarity.

1. A transition *τ* from *A ϵ σ* to *B ϵ* ∑ can be written (*A, B*) or (A →B).
2. *τ* = (A → B) is *defined* for *A* and *leads to B*, written *A ≺ τ* and *τ ≻B* respectively. The notation is transitive to bundles and sets, i.e. *A ≺ π ⇔ ∃ τ ϵ π,A ≺ τ* and *π ≻B ⇔ ∃ τ ϵ π τ ≻B,* respectively.
3. A bundle *π* forms a tuple (*S, T*) with start and target symbols S ={ *σ|σ ≺ τ,τ ϵ π*} and *T* = {*σ |τ ≻σ*, *τ ϵ π*},respectively.
4. If a transition bundle *π*_*i*_ is *true*, then so are all contained transitions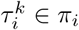

Transitions form terms which are independent of symbols. Consider the symbol *A* and the transition A→B, both of which are propositional terms, in the term A Λ (A→B) If *A* is *true*, then it can be deduced logically that *B* is also *true*, written A Λ (A→B) ⇒B. Here, Λ is the *logical and* operator. *A* forms a *precondition* for the transition (A→B). As *A* is true, the precondition is met and thus also the transition is true. *B* is the *conclusion* of the entire term. Order of evaluation is not specified during logical deduction. Therefore, sequential evaluation of transitions is made explicit as follows.

**Definition 3** (Transition evaluation). *A configuration* Ω *ϵ P*(∑) *of an MTS ℳ is* evaluated *according to the functions*

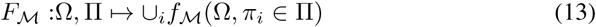

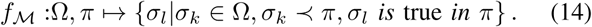

The union in Equation 13 goes over all transition bundles contained in. Hence, evaluation yields all symbols which are true given an initial configuration. The function *F*_*ℳ*_(ΩII,) allows recursive usage for evaluation until a target symbol is reached. Consider the following example with four symbols *A, B, C, D*, where *A* is the start and *B* the target. Hence the initial configuration is Ω = *{A}*. Furthermore, the following transition set, bundles, and points are defined.

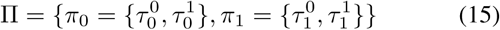

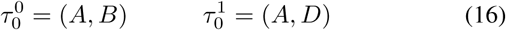

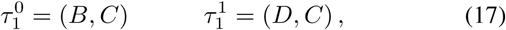

It requires two recursive evaluations, i.e. *F*_*ℳ*_(*F*_*ℳ*_(Ω,II),II), until the target symbol is found. This is similar to the evaluation of transition functions in RL [120]. However, the set notation allows the superposition of several symbols at the same time.

### C. Encoding capacity of the Universal MTS

The definition of *π* introduced a *bundling trick* which provides several benefits to analyze the computational logic and storage requirements of an MTS, especially when viewed in the light of neural encodings. Consider the following thought-experiment. Suppose that the generation of a bundle (e.g. a neuron) is energetically expensive, however the addition of a transition point (e.g. a dendritic spine) to an existing bundle is comparably cheap. To avoid evolutionary pressure [96], the goal is thus to minimize the overall cost. Optimizing this cost corresponds to maximizing the number of transition points while minimizing the number of bundles. As will be shown now, it is not possible to merge arbitrary transition points in one bundle without violating the constraints of MTT.

**Theorem 1.** *Let* σ ϵ ∑, *ℳ an MTS on the alphabet* σ,II *the corresponding transition set, and π* = (*S, T*) *a transition bundle. ℳ generates* valid *sequences if and only if the following conditions hold.*

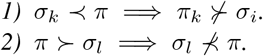

*Proof.* 1) From Axiom 1 it follows immediately that any transition *π* which is defined for *σ*_*k*_ and leads to *σ*_*k*_ violates the non-stationarity condition. 2) Without loss of generality, consider the three symbols *σ*_0_,*σ*_1_,*σ*_2_ *ϵ* ∑ and *σ*_0_,*→σ*_1_,*→σ*_2_ but *σ*_0_ ↛ *σ*_2_. This yields the transition points *τ*_0_ = (*σ*_0_, *σ*_1_) and *τ*_1_ = (*σ*_1_,*σ*_2_). Assume further that *τ*_0_ and *τ*_1_ are bundled in *π*, and that *σ*_0_ and *π*_1_; are *true*. It follows that *σ*_0_Λ*τ*_0_⇒ *σ*_1_ However, *σ* _1_Λ*τ*_1_ ⇒ *σ* _2_ and thus *σ* _1_Λ *π* ⇒ *σ*_2_ This contradicts the assumption and violates the coherency constraint.

Therefore, the sets of symbols *S, T* for a *π* = (*S, T*) have to be mutually exclusive, i.e. S ∩ T = ϕ

**Definition 4** (Minimality, universality). *An MTS ℳ is* minimal *if there exists only one π*_*i*_ *for any σ*_*k*_, *i.e σ*_*k*_ *≺ π*_*i*_ *⇒ σ*_*k*_ *⊀ π*_*j*_ *for any j ≠ i. In a* universal *ℳ, any arbitrary transition between two symbols σ*_*k*_, *σ*_*l*_ *is possible.*

**Corollary 1**. *The input set S*_*i*_ *of a transition bundle π*_*i*_ *is singleton for a* minimal *and* universal *ℳ.*

*Proof σ*_*k*_ *≺ π*_*i*_ and *π*_*i*_ ≻ *σ*_*l*_, ∀ *l ≠k* According to Theorem 1, *σ*_*l*_*⊀ π*_*i*_, ∀ *l* ≠ *k*

**Corollary 2.** *Let* Σ *be an alphabet of size M, II a transition set of N transition bundles π*_*i*_ = *{S*_*i*_, *T*_*i*_*} for a minimal universal M. When all possible transitions are realized, then M* = *N. Proof.* Construct the graph *G* of *M* in which each transition is represented by a node, and any symbol by a directed edge. For example, consider the three symbols *σ*_0_, *σ*_1_, *σ*_2_ which are connected by the transitions *τ*_0_ = (*σ*_0_, *σ*_1_) and *τ*_1_ = (*σ*_1_, *σ*_2_). The corresponding graph has only two nodes for *τ*_0_ and *τ*_1_ which are connected by a single directed edge representing *σ*_1_.The directed graph for a complete *universal minimal M* in which all transitions are realized is fully connected. Reducing any pair of directed edges to a single undirected edge yields a fully connected undirected graph. According to Theorem 1, *S*_*i*_ *\ T*_*i*_ = ɸ for any *π*_*i*_. Therefore, only those transitions can be bundled which are not connected by an edge in *G*. Furthermore, the number of independent nodes in *G* is equivalent to the chromatic number of the graph, i.e. the minimal number of colors which can be assigned to nodes of a graph, known as graph-coloring problem [121]. The chromatic number of a fully-connected graph equals the number of nodes. Conclusively, *M* = *N*.

An example of the reduced graph for a sequence of four symbols *σ*_1_,…, *σ*_4_ is depicted in Figure 7b. The figure shows transitions as vertices, and edges correspond to symbols. Removing the directedness of the graph by fusing two symbols means that each edge is associated with two symbols.

### D. Brief discussion of the Universal MTS with implications for neural networks

Following Corollary 2, an implementation of a universal minimal *ℳ* requires to have as many entities to store transition bundles as it has symbols. Furthermore, these encoders have to decorrelate from their target symbols to fulfill Theorem 1.

Consider a neural implementation in which a neuron represents a transition bundle. To make efficient use of the neuron, it has to represent multiple transitions and thereby expose several receptive fields. However, each transition also dictates that the neuron has to dissociate from target symbols. Combined, each receptive field of such a transition neuron is suggested to consist of an on-region in which it associates to the symbol for which it is defined, and an off-region in which it decorrelates. In addition, the transition system requires the possibility of representing a *logical and* operation, feasible in neural networks [123]. Finally, each neural transition encoder will co-activate with any symbol for which it is defined during recursive retrieval.

### E. Multi-Transition Systems in Euclidean space

The space which is constructed by symbols *δ*_*i*_ and transitions *τ*_*j*_ above is the discrete topological space with the induced discrete metric. However, the world in which animals reside is not discrete and arbitrary jumps between two locations are infeasible. In particular, the perceived environment corresponds to a complete metric space, i.e. the Euclidean, from now on simply called *metric space*. Hence, an MTS *ℒ* which encodes transitions in a metric space has different constraints than a universal MTS *ℳ*

Encoding transitions between locations in a metric space depends on the detection of two consecutive positions. The following analysis is based on the assumption that there exists a continuous signal which depends on and uniquely identifies each possible location of the animal. In terms of the Euclidean space *M*, this corresponds to locations x *ϵ M*. Certainly an animal does not have access to coordinates. However, other stimuli are likely to provide the necessary information. For instance, geometrical information combined with head direction signals is sufficient to represent singular locations, which was demonstrated in the BV cell model presented by Barry et al. [18].

According to Definition 1, an alphabet is finite. This can be understood to correspond to a finite number of neurons which have to represent locations. However, the alphabet Δ of *spatial* symbols *δ*_*i*_ has to represent the continuous signals x of the input space *M*. This corresponds to the well-known *sampling theorem*.

**Definition 5** (Spatial symbol, enablement, and assignment). *Let δ*_*i*_ *ϵ* Δ *be spatial symbols according to a sampling process of a complete metric space D* = (*M, d*). *Each δ*_*i*_ *is centered at a* x_*i*_ *ϵ M.*

*A point* p *ϵ M* enables *δ*_*i*_ *if it is within the* support *of δ*_*i*_ *given by the open ball B*_*i,s*_ *of radius r*_*s*_, i.e. *B*_*i,s*_ = *{*p *ϵ M|d*(x_*i*_, p) < *r*_*s*_*}.*

*The point* p *is* assigned *to the closest δ*_*i*_, *i.e. δ*_*i*_ *for which d*(x_i_, p) *is minimal. Given two adjacent symbols δ*_*i*_, *δ*_*j*_, *then r*_*ω*_ = *||d*(x_*i*_, x_*j*_)*||/*2, *describing a ball B*_*i, ω*_ *of radius r* _*ω*_.

The definitions of enablement and assignment can be interpreted in the following way. The region in which a spatial symbol is enabled can be understood as its receptive field, and according to the definition, multiple spatial symbols can have overlapping receptive fields. In contrast, assignment identifies the closest symbol, for instance as a result of a winner-take-all mechanism.

According to the Petersen-Middleton theorem [122], the ideal sampling strategy for two-dimensional continuous signals and therefore placement of spatial symbols *δ*_*i*_ is a hexagonal arrangement. The sampling process can also be understood as a solution to the problem of packing spheres with diameter *r*_*ω*_ as densely as possible. The sphere packing problem also yields a hexagonal lattice in the two dimensional case [124], [125].

Assuming an ideal sampling process, the question remains about the optimal distribution of transition bundles.

Theorem 2. *Let D* = (*M, d*) *be an Euclidean space. Let ℒ* = (*P*(Δ),Г) *be a minimal transition system on D such that the countably finite alphabet Δ corresponds to the densest optimal covering with respect to r*_*ω*_.

1. *The number of transition bundles γ*_*i*_ *ϵ* Г *is constant.*
2. *2) The occurrence of any transition bundle γ*_*i*_ *is periodic.*

The theorem is proved by its corresponding graph-coloring problem, introduced in Subsection IV-C.

*Proof.* The densest arrangement of spatial symbols according to the Petersen-Middleton theorem is a hexagonal lattice [122]. Furthermore, transitions between symbols are only possible between adjacent symbols. Consequently, the corresponding transition graph is not complete, i.e. only neighboring transitions are connected. The chromatic number of the resulting graph is 3 and the occurrence of colors is periodic.

An example for the two dimensional arrangement of symbols is depicted in Figure 7c, and one solution of the graph coloring problem in Figure 7d.

As mentioned above, likely candidates to provide an input space which yields unique signatures for arbitrary locations in confined environments are BV and HD cells. However, an analysis which uses biologically plausible afferents is postponed to future work.

## V. DISCUSSION

A biological plausible self-organizing model of GCs, proposed to encode transitions, was derived in Section II. Hexagonal grid fields emerged due to recurrent dynamics and to optimally encode transitions. Subsequently, Section III introduced a scale-space model for GCs to remedy the behaviorally problematic runtime of spatial transition systems during path planning. Finally, MTT was introduced in Section IV to mathematically examine the storage requirements both for episodic as well as spatial transitions.

Learning sequences and transitions in the Hippocampus was explored previously [126], [127]. However, these studies ignored spatial information or GCs. Spatially modulated inputs and association of motor commands and rewards to spatial transitions were already suggested by Cuperlier et al. [110], [111]. In their model, places and transitions are stored separately. In addition, their model contains biologically plausible interactions between the motor cortex, and was demonstrated in a real-world example using a robot. Hirel et al. extended the model [112] to incorporate reinforcement learning for the acquisition of goal-directed transitions in a biologically plausible manner. However, these models did not differentiate between spatial and temporal transition systems. Furthermore, sequences were not defined rigorously and optimality of transition encoding was not subject of the studies.

The work presented here used dendritic computation for the self-organization of GCs. A similar model was suggested previously by Kerdels et al. [128]. However, the authors proposed that GCs perform sampling of a circular input space for the purpose of localization. Thereby, the cells perform Voronoi clustering, which yields a hexagonal arrangement of grid fields in the ideal case. In addition, the model presented in Section II is similar to the work by Si et al. [129] and Kerdels et al. [128] in that it suggests that GCs of one module properly align to each other due to recurrent dynamics.

### A. Biological interpretation of MTT

For arbitrary sequences and transitions from one symbol to another, the mathematical proof shows that as many transition encoders are required as there are symbols. In Euclidean metric space, the result differs in that only a finite number of transition encoders is required given the assumption of unique spatial input. Widloski et al. recently proposed a model of GC learning which exhibits similarity with respect to the expected behavior of the cells [130]. The authors demonstrated that a spiking neural network converges to a hexagonal arrangement of responses given spatially unique inputs.

Retrieval of sequences from an MTS requires both symbols and transition encoders. Neurons which represent transitions between spatial symbols will thus co-activate with and express similar place field behavior as the neurons encoding the symbols. Likely candidates for the storage of symbols and transitions appear to be PCs of CA1 and CA3. However, it is unclear if episodic transitions are stored within recurrent collaterals, plenty of which were found for CA3 [73], [131], or if there exists a suitable recurrence from CA1 to CA3. Another possibility is an embedding in the trisynaptic loop, which spans CA1, CA3, and EC. For the first case, disruption of activity within CA1 *or* CA3 should lead to a decline of performance in tasks in which temporally accurate sequences are required.

It is expected that the deliberately abstract MTT can be applied to other domains which require transitions between symbols.

### B. A note on spatially unique input

All three introduced aspects, i.e. MTT, the self-organizing GC model, as well as the scale-space model, were developed under the assumption of spatially unique sensory inputs. Although the models used coordinates, it is proposed that the BV space forms a suitable and biologically plausible sensory representation. This is exemplified by the BV model by Barry et al. [18], who demonstrated that afferents from BV cells are sufficient to plausibly generate and correctly predict the firing of PCs with respect to the environment. Consequently, it is assumed that the coordinates can be replaced by a similar presynaptic sensory state. In fact, it is expected that coupling GCs and BV cells in the scale-space model will be able to explain recent findings in which GCs appeared to be influenced by the geometrical layout of the environment [132], [133].

### C. Remarks on the self-organizing model of grid cells

The self-organizing model of Section II quickly develops cells with a gridness score above zero for realistic trajectories of rodents. Furthermore, the dynamics align the orientation of all participating cells of one grid module. Competitive dynamics were required to achieve co-orientation of several cells, which is supported by Couey et al. who found that recurrent connectivity in the mEC is primarily inhibitory [84].

The model uses an animal’s speed to adapt learning of spatial transitions. In particular, the rule defined in Equation 10 modulates the learning rate inversely proportional to the running speed of the animal. Hence, knowledge about the linear speed of the animal is required, which was recently discovered to be represented by speed cells in the mEC [134]. The effect is expected to be realized in either of the two ways. On the one hand, inhibitory interneurons could suppress the activity of the currently spiking GCs more strongly the faster the animal runs. On the other hand, a purely excitatory mechanism could be based on influencing GCs which sample future sensory states. Due to the findings of Couey et al. [84], the first mechanism appears to be more likely.

The presented model is independent of movement direction. On the contrary, the model depends on the instantaneous perception of the environment and identification of locations, similar to other previously proposed self-organizing models [88], [89], [128]. However, it is expected that activity of simulated GCs will slightly follow the head direction of an animal when the visual system is modeled accurately. There are two reasons for this expectation. First, the visual system of rodents is slightly forward facing, making it more likely that locations are recognized that are in the viewing direction. Second, one of the main purposes of the transition system is to plan trajectories towards goals. It appears to be more relevant to plan towards where an animal is facing, rather than were it is moving. In principle, this could explain a recent statistical analysis by Raudies et al. [57], who showed that GCs activity correlates more strongly with head direction than with movement direction.

The presented self-organizing model is limited due to negligence of temporal dynamics on the synaptic level. For instance, the receptive fields with on-center and off-surround dynamics are expected to be a result of temporal interactions between presynaptic afferents in combination with an asymmetric STDP learning rule. Furthermore, the model was only shown for three cells which are likely to organize themselves hexagonally and evenly distribute across the input. However, stochastic input in combination with a suitable STDP rule is expected to generate hexagonal arrangements and GCs with partially overlapping receptive fields. Finally, the programmatic global and local winner-take-all mechanisms are expected to also be remedied by a detailed model with accurate temporal dynamics.

### D. Interpretation of the scale-space model and related areas

The emerging scale-space structure of Section III can be interpreted in the following way. The finest resolution of the spatial transition system reacts to transitions on the perceivable level, represented by presynaptic activity. In other words, detection of spatial locations during the dendritic computations corresponds to an *identity* function. The target region of any transition which is detected on this scale corresponds to any location with which an animal can interact directly. Larger scales allow the animal to asses viability of movement towards locations which are further apart. To allow comparison of such displaced locations, the corresponding sensory representations are low-pass filtered in discrete steps. In turn, these comparisons allow to perform approximate look-ahead. Look-ahead was already proposed to be performed in the entorhinal-hippocampal loop by Kubie et al. [135]. However, the authors did not address optimality of the encoding.

During learning, the simplified scale-space model requires co-activation of both spatially modulated afferents, as well as already acquired spatial symbols. The reason is that on the one hand, GCs are assumed to detect spatial transitions directly on the input space due to dendritic computations. On the other hand, these transitions must be related to already learned PCs. However, sensory information may not be available during retrieval, especially mental travel. Hence, retrieval operates only due to activated PCs, which requires to drive GC activity only on behalf of PCs. It is therefore expected that these operations are implemented hetero-synaptically in such a way that there is a clear distinction between learning and retrieval in the mEC.

The described recursive technique to construct multi-scale look-ahead transition encoders generates a Gaussian pyramid, or scale-space, well known in the computer vision and signal processing communities [136]–[138]. Among others, scale-spaces were used to describe biologically plausible retinal and visuocortical receptive fields [139]–[141], and it was proved that a scale increment of 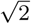 is optimal for Gaussian functions [138]. In image processing, application of a Gaussian pyramid corresponds to consecutively smoothing an image, i.e. an input gets low-pass filtered by which fine-scale information is removed. Finally, this allows to detect features in the input across several scales [142], [143], thereby making feature detection scale invariant. The same principle is used in the scale-space model for GCs to search for feasible transitions on larger distances. Essentially, comparison of two distal locations for look-ahead requires to ignore fine structures.

### E. Temporal buffering and a relation to Theta phase precession

Lindeberg points out that access to temporally buffered information is necessary to construct scale-space structures [144]. Certainly, this was also required for the construction of the scale space in Section III. The temporal buffer was required to bind large-scale transition encoders to consecutive places across long distances. Hence, GCs of larger scales require excitation not only from the current sensory perception, but also from PCs which encode past and future locations on a linear track. This observation is supported by findings which report that excitatory drive from the Hippocampus is required for grid cells [54]. The observation predicts that the temporal integration window of GCs from larger scales needs to be increased when compared to cells from smaller scales. Or, that some other mechanism exists which prevents GCs of larger scales from firing before the necessary afferents from PCs were accumulated. Most importantly, it requires a biologically plausible mechanism to buffer data temporally.

The most likely candidate for a temporal buffer is Theta phase precession [34], [35]. When an animal is running, several PCs which correspond to past locations, the PCs of the current location, as well as PCs which encode future locations spike *in order* of their traversal within one Theta cycle. The reported maximal compression ratio is of the order of 10: 1 [33],i.e. at most ten consecutive places are represented within one theta cycle. One iteration of the proposed transition system requires at least two neural memories. Given approximate numbers of neural activation and axonal delay of 5 – 10 ms, one iteration requires 15 ms on average. In concordance to the reported compression ratio, at most 10 iterations of the transition system fit into one Theta cycle, which oscillates between 6–10 Hz [145]. Conclusively, Theta is proposed to form a main-loop with nested sub-loops that iterate the transition system. Without other mechanisms, it is estimated that on average 5-7 stable grid scales can be formed on top of this temporal buffer, depending on the frequency of Theta. It is expected that temporal compression during SPW-R contributes to the formation of grid scales, which may lead to larger scales.

### F. Links to computer science and robotics

Skip lists [114], used as inspiration for the look-ahead in the scale-space model, operate on one-dimensional lists of data. Several related data structures which form hierarchies for two dimensional data are known in computer science. For instance, quad-and octrees subdivide dimensions of an input space to accelerate queries exponentially [146], [147]. On topological data without global coordinates, contraction hierarchies were shown to significantly improve search time [148], [149]. The scale-space model of GCs demonstrates how these data structures can be extended to probabilistic data.

An MTS creates a topological representation of perceived locations. Dabaghian et al. already suggested that the rodent Hippocampus forms a topological map [25], [150]. However, the novel scale-space model not only links immediately neighboring locations but allows to extract approximate relations between remote places. Retrieval of a path according to the scale-space model corresponds to the well known Dijkstra’s algorithm [151], or more specifically to A* [152]. The latter algorithm was developed for robotic navigation and is an extension of the first. A* uses heuristic information to accelerate search of viable trajectories, similarly to look-ahead transitions in the scale-space model. The benefit of a topological map, formed on spatially modulated sensor data and augmented with look-ahead, is the avoidance of a global coordinate system. It is thus expected to be useful in indoor robotics systems without access to global coordinates.

### G. Future work

Several simplifications and abstractions were necessary to reduce the complexity of the introduced models. An implementation which is currently in development will use a spiking neural network to address the issue of missing temporal dynamics, absent in the self-organizing model of Section II. Early results of the model show that on-center and off-surround fields for dendritic computation form as expected, and that several cells exhibit partially overlapping grid-like fields.

Furthermore, an extended model of Section III is in development which includes RL as a mechanism for trajectory selection. The model presented in Section III lacks a quality measure to narrow down multiple possible trajectories to a singular best fit and only reports the existence of any viable path. It is expected that the previous work by Hirel et al. [112] can be adapted and reproduced. Subsequently, the RL mechanism will be combined with the spiking neuron model. It is expected that the observation of pre-play of trajectories, reported by Pfeiffer et al. [31], can be reproduced in the combined model.

## VI. CONCLUSION

This work proposed an entirely novel function of grid cells,i.e. optimal encoding of transitions across multiple scales in form of a multi-transition system. First a biologically plausible model of grid cells in a competitive network was developed. Then, the spatial multi-transition system was extended into a scale-space model for transition encoding to solve behaviorally problematic run-times. Given the assumption of probabilistic detection of locations, it was shown that the optimal scale increment for transition encoding in the scale-space model is *√*2. Finally, Multi-Transition Theory was introduced to formally analyze transition systems. It was shown that a hexagonal arrangement of transition encoders minimizes the number of required encoders in two-dimensional Euclidean space. The results give rise to future developments. Extended models with spiking dynamics are in development to address biologically plausible reinforcement learning for trajectory selection and input space representation. Furthermore, the results are currently used for the development of novel mobile swarm-robotic systems which operate on and communicate with probabilistic data similar to the principles of neural systems.Conclusively, this work addressed spatial navigation on all three levels of analysis suggested by David Marr [153]. 1) The purpose of the computation, which is twofold. On the one hand an animal requires to record locations. On the other hand, the recorded locations have to be recalled in such a way that trajectory planning can be achieved in a behaviorally acceptable run-time. 2) The algorithmic description and especially the scale-space model were a result of the computational purpose and its behavioral relevance. 3) A biologically plausible implementation for GCs as well as the scale-space multi-transition system were demonstrated.

## VII. ACKNOWLEDGMENTS

The author would like to thank Christoph Richter and Jörg Conradt for invaluable discussions and feedback during the research, as well as their support and suggestions on the manuscript.

